# STCS: A Platform-Agnostic Framework for Cell-Level Reconstruction in Sequencing-Based Spatial Transcriptomics

**DOI:** 10.64898/2026.02.26.708370

**Authors:** Lixia Chen Wu, Xinyu Hu, Fengwei Zhan, Chuhanwen Sun, Jose Gonzales, Rachel Ofer, Tyler Tran, Michael P. Verzi, Liping Liu, Jiekun Yang

## Abstract

Sequencing-based spatial transcriptomics technologies, including Visium HD and Stereo-seq, now enable transcriptome-wide profiling at subcellular resolution. However, these platforms generate measurements over spatially barcoded units rather than biologically segmented cells, creating a fundamental bottleneck for cell-centric analysis and interpretation. Robust reconstruction of coherent single-cell transcriptomes from high-density spatial bins remains an unresolved computational challenge.

Here we present **STCS (Spatial Transcriptomics Cell Segmentation)**, a platform-agnostic framework that reconstructs cell-level gene expression profiles by integrating nuclei segmentation with a joint transcriptomic–spatial distance model. STCS is governed by two interpretable parameters and incorporates a reference-free parameter selection strategy based on internal stability and spatial coherence metrics, enabling adaptable deployment across tissue types and technologies without requiring matched ground-truth annotations.

We benchmark STCS on a Visium HD human lung cancer dataset with matched Xenium-derived cell segmentation, enabling direct cell-level validation, and on high-resolution Stereo-seq mouse brain data to assess cross-platform generalizability. Across multiple evaluation dimensions—including cell-type agreement, spatial organization, gene-expression fidelity, and compositional accuracy—STCS achieves consistent improvements over existing methods while preserving biologically coherent spatial structure.

As sequencing-based spatial transcriptomics is rapidly adopted across biomedical research, STCS provides a broadly applicable and open-source solution for reconstructing cell-resolved transcriptomes, facilitating more reliable downstream analyses and cross-platform integration.

## Introduction

Spatial transcriptomics (ST) has emerged as a transformative technology that enables high-throughput gene expression profiling while preserving the spatial organization of transcripts within intact tissue sections (1). It has rapidly expanded into a diverse family of experimental platforms that differ in spatial resolution, molecular capture strategies, and sequencing throughput, collectively enabling increasingly detailed interrogation of spatial gene regulation in complex tissues (2; 3).

Most sequencing-based spatial transcriptomics platforms achieve transcriptome-wide coverage by partitioning tissue sections into dense grids of barcoded spatial units, commonly referred to as spots or bins, which capture transcripts originating from micrometer-scale regions of tissue (4; 5). Visium HD is one such platform, profiling gene expression using subcellular barcoding at approximately 2 µm resolution, with subcellular barcodes subsequently aggregated into larger spatial units, typically aggregated in 8 µm for downstream analysis and annotation (6). Similarly, Stereo-seq generates ultra-high-density spatial transcriptomics data using DNA nanoball arrays with submicrometer feature spacing on the order of 200 nm (7).

Despite their high spatial density, sequencing-based spatial transcriptomics platforms generate measurements defined over spatially barcoded units rather than over biologically segmented cells (1; 8). While subcellular or nanoscale barcoding increases spatial resolution, it does not define cellular boundaries or assign transcripts to individual cells. Consequently, expression profiles derived from spatial bins represent aggregated transcriptional signals rather than coherent single-cell transcriptomes.

Imaging-based spatial technologies such as Xenium partially address this limitation by detecting individual RNA molecules in situ through cyclic fluorescence imaging and assigning them to segmented cells based on their cell boundary protein staining whenever possible (9; 10). However, these approaches rely on predefined gene panels, restricting transcriptome coverage and can exclude genes important for unbiased cell-state characterization and discovery-driven analyses (11).

Many downstream analyses in spatial transcriptomics are inherently cell-centric, requiring gene expression measurements to be defined at the level of individual cells (8). However, current sequencing-based spatial transcriptomics platforms provide genome-wide expression profiles without intrinsic cellular resolution, whereas existing cell-resolved spatial technologies rely on predefined targeted gene panels and are consequently constrained in transcriptome breadth and scalability (1; 11)

Several methods have been developed for cell segmentation on Visium HD data. Bin2cell uses StarDist (12) to segment nuclei and expands them by a fixed distance to approximate cell boundaries, assigning overlapping bins based on genetic similarity (13). However, this approach may generate cells with spatially disconnected bins, and fixed-distance expansion can lead to inaccurate boundary estimation and over extension. More recently, 10x Genomics released an updated version of Space Ranger (14) with Visium HD cell segmentation functionality, but it is not open source yet. Here, we introduce STCS, a computational framework that reconstructs cell-level units by aggregating bins into coherent cellular representations, which enable downstream analyses, including cell-type annotation using marker-based approaches or automated classifiers. Compared with existing approaches, STCS achieves improved cell boundary coherence and competitive performance across cell-type accuracy, cell-type-specific gene expression and spatial consistency metrics. Importantly, STCS is fully open-source and designed to be platform-agnostic, enabling application across diverse sequencing-based spatial technologies.

In addition to supporting Visium HD-scale data, STCS operates on any spatial transcriptomics dataset in which transcriptional measurements are spatially aligned with tissue histology images. Despite the substantially smaller spatial units and higher measurement density of Stereo-seq relative to Visium HD, STCS remains effective in reconstructing coherent cell-level expression profiles from nanoscale spatial data.

## Results

### STCS enables self-defined parameters for accurate prediction on different slides

STCS first processes H&E images using StarDist to obtain nuclei segmentation results. For bins located outside any nucleus, neighboring nuclei within a fixed search radius are identified as candidates. STCS then calculates a combined distance for each candidate, integrating genetic distance and weighted spatial distance. Each bin is assigned to the nearest nucleus based on this combined distance, generating single-cell level results. This assignment process is governed by two key parameters: the search radius (*S*) and the spatial weighting factor *λ* (Fig. 1A). The search radius defines the maximum spatial range for candidate nucleus selection, thereby influencing the spatial extent of reconstructed cells. The parameter *λ* modulates the relative contribution of spatial distance in the combined distance metric, regulating the balance between transcriptomic similarity and spatial proximity.

**Figure 1:**
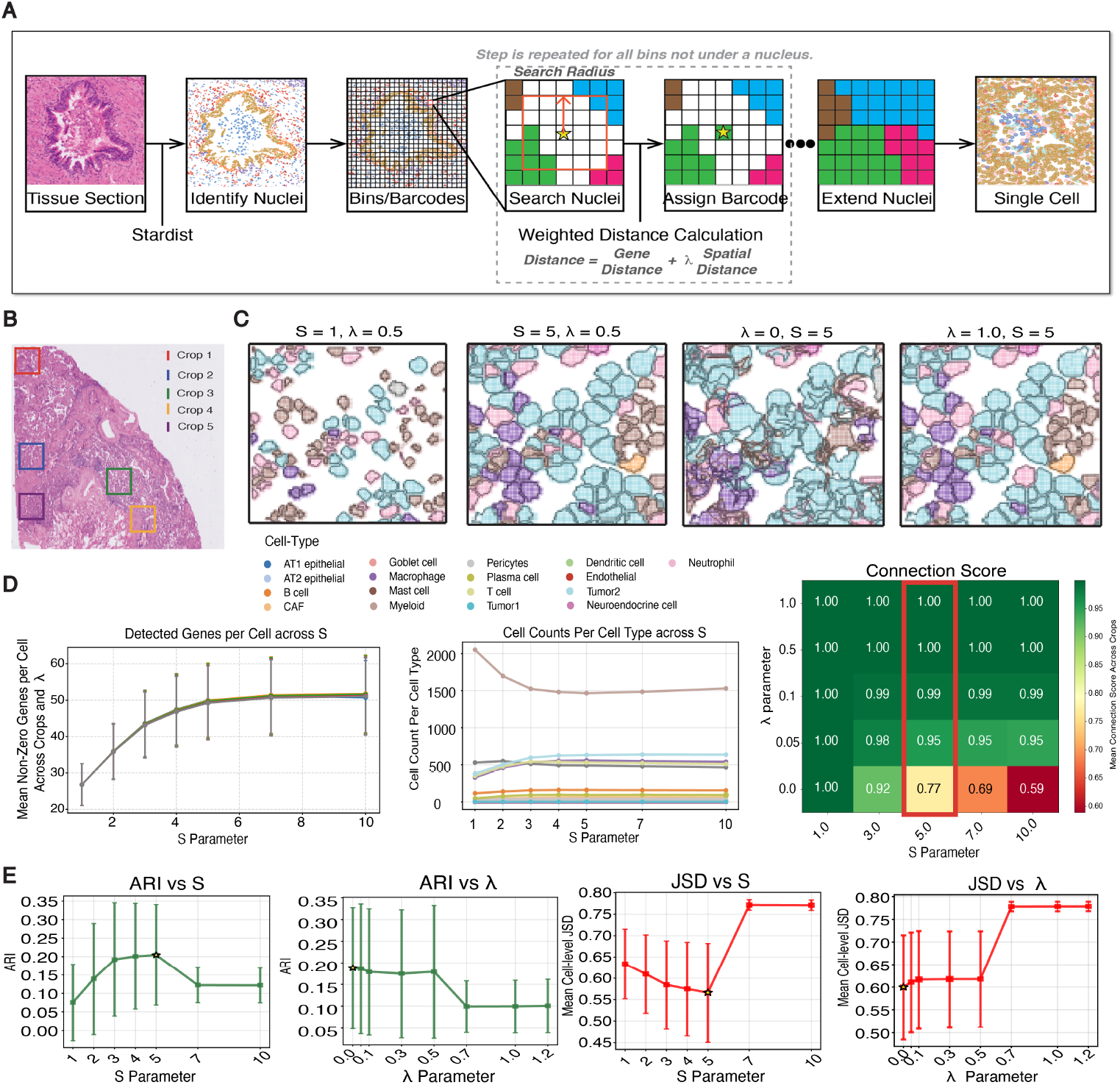
Pipeline and parameter tuning of STCS. **A)** Overview of STCS pipeline. **B)** Five randomly selected crops used for parameter evaluation. Parameter tuning was performed over combinations of search radius S and spatial-weight parameter *λ*. **C)** Cropped ROI illustrating STCS assignments under different S, *λ* combinations. **D)** Parameter selection metrics computed across the five crops shown in (B). Left: Mean detected genes per cell across S. Line plot with standard deviation bars and each connected center point represents the mean number of detected genes per cell across *λ* values, illustrating the dependence of gene detection on the search radius. Center: Cell counts per cell type across S. Lines represent individual cell types, and counts correspond to the mean values across all *λ*, representing the stability of cell-type prediction. Right: Connection score across parameter combinations (S,*λ*). This metric quantifies the spatial coherence of assigned cells, where higher values indicate more contiguous and structurally consistent cell assignments. **E)** Adjusted Rand Index (ARI) and Jensen–Shannon Divergence (JSD) are shown versus S (mean over all *λ*) and versus *λ* (mean over all S). Stars denote the best mean value and error bars represent standard deviation. These metrics were computed by comparing STCS-segmented cells across five crops to matched Xenium cells, with results averaged across crops to assess performance across different S and *λ*.

To characterize parameter sensitivity, five randomly selected regions of interest (ROIs) were sampled from the human lung cancer slide (Fig. 1B), and STCS was executed across a range of search radius and spatial weighting combinations (Fig. 1C). Qualitative comparisons demonstrate that increasing the search radius, while holding *λ* constant, systematically enlarges reconstructed cellular domains, reflecting expansion of the candidate assignment neighborhood. In contrast, variation of *λ* primarily affects the spatial structure of assignments: larger values promote spatially contiguous bin aggregation, whereas smaller values permit increased spatial fragmentation. These patterns confirm the distinct functional roles of *S* and *λ* in governing reconstruction scale and spatial coherence.

To guide parameter selection, we systematically evaluated performance across multiple quantitative criteria (Fig. 1D). During search-radius optimization, smaller values of S resulted in reduced numbers of detected genes per cell, consistent with constrained spatial aggregation. The mean detected-gene count increased progressively with S but reached a pronounced plateau at *S* ≥ 5, indicating saturation of transcript capture within reconstructed cellular domains. Beyond this threshold, further increases in S produced negligible changes, suggesting that additional neighborhood expansion does not influence cell-level transcriptional profiles.

This stabilization pattern was further supported by cell-count analysis per cell type, where cell identities were assigned using the CellTypist model (15) for each crops. The number of cells assigned to each predicted cell type remained largely invariant once S exceeded 5 (Fig. 1D). Collectively, these observations indicate that *S* = 5 is sufficient to capture stable transcriptomic structure and cell-type prediction properties while avoiding unnecessary computational overhead.

For spatial-weighting calibration, we assessed structural coherence using the connection score, which quantifies the proportion of bins forming spatially contiguous regions within reconstructed cells. Lower scores reflect fragmented spatial assignments, whereas values approaching unity indicate coherent cellular structures. Connection scores increased monotonically with *λ* and stabilized near *λ* = 0.5 (Fig. 1D). On the basis of these results, we selected *λ* = 0.5 and *S* = 5 for this data set.

We further validated the selected parameter configuration using Xenium annotations from the same slide. STCS-predicted cells were aligned to Xenium cells based on nuclear spatial overlap, enabling direct cell-level comparisons. Agreement between reconstructed and reference annotations was quantified using the Adjusted Rand Index (ARI) and Jensen–Shannon divergence (JSD, Fig. 1E).

Both ARI and JSD exhibited parameter-dependent behavior across *S* and *λ*. ARI values peaked near *S* = 5 and *λ* = 0.5, followed by a systematic decline as either parameter increased. This reduction in ARI indicates diminished agreement with reference cell identities. JSD exhibited the complementary behavior expected for a divergence metric. Divergence was minimized near *S* = 5 and *λ* = 0.5 and increased with further parameter escalation, reflecting progressively greater deviation between predicted and reference cell-type distributions. These analyses yielded consistent conclusions: optimal performance was observed at *S* = 5 with *λ* = 0.0 or 0.5.

These concordant trends between internal quality metrics and external validation measures demonstrate that the parameter configuration identified through unsupervised evaluation of randomly sampled ROIs closely aligns with the optimum determined by ground-truth comparison. This agreement establishes a computationally efficient, reference-free parameter selection process for optimal STCS configurations.

### STCS showed a slight improvement over existing methods on a Visium HD slide with ground-truth cell segmentation

After determining the optimal parameters using image crops, we applied the selected parameter set to the entire slide. Cell-type annotations were generated using CellTypist trained on a human lung cancer single-cell reference dataset (16). The resulting predictions showed strong qualitative concordance between the Xenium ground truth and STCS segmentations (Fig. 2A).

**Figure 2:**
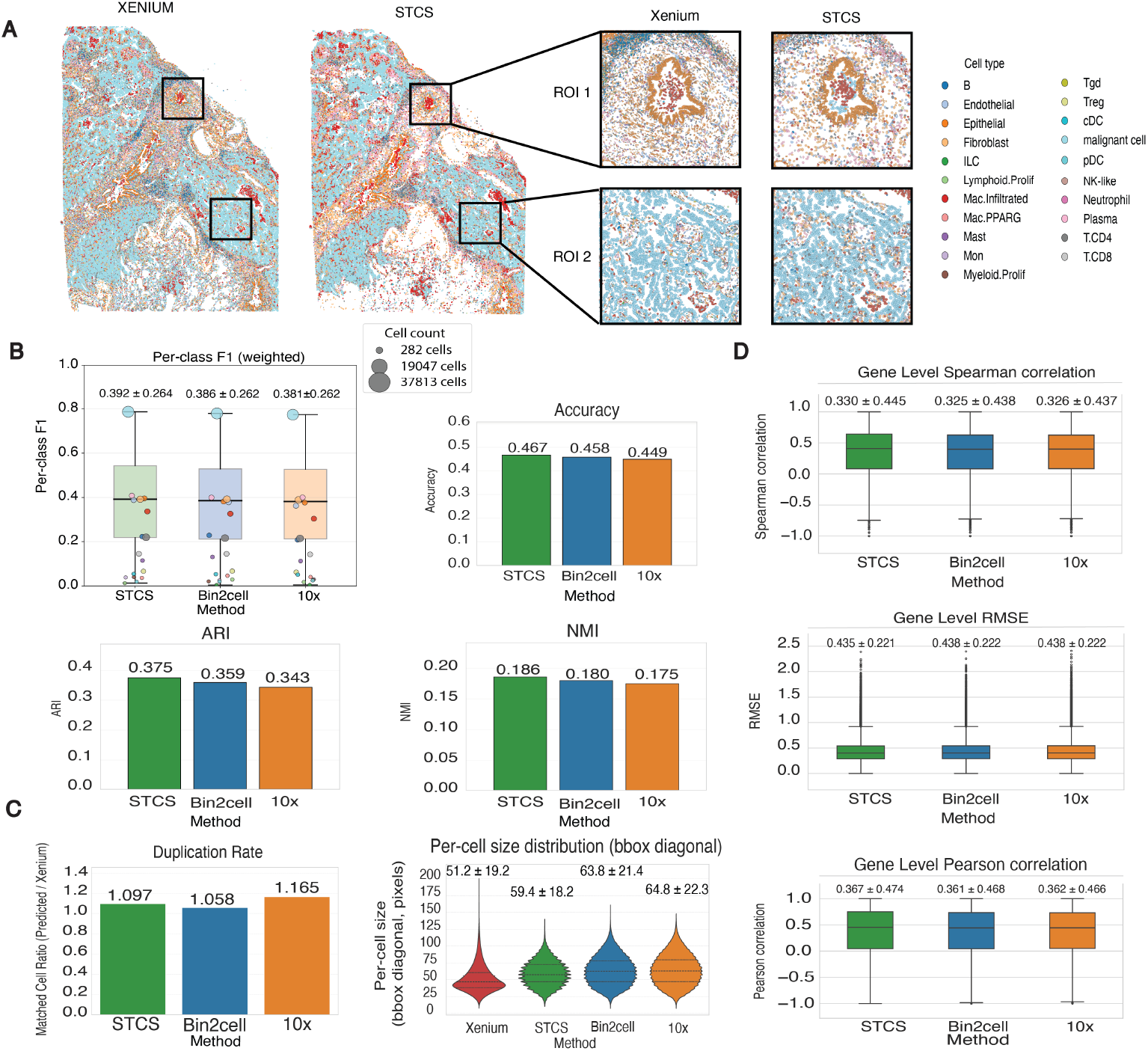
Metrics comparing STCS with existing methods and Xenium-derived ground-truth cell segmentation. **A)** Whole-slide cell segmentation results and ROI crops, colored by assigned cell types. Cell-type labels for all methods and the Xenium ground truth were obtained using a trained CellTypist model. **B)** The four metrics summarize cell-level cell-type agreement between reconstructed cells and Xenium ground truth, computed on matched Xenium–prediction pairs and compared across STCS, Bin2Cell, and 10x Space Ranger segmentations. Upper left: Weighted per-class F1 scores. Dots represent per-cell-type F1 values, colored by the cell-type annotations shown in (A) and scaled by class support (number of matched cells). Boxes depict the support-weighted median and interquartile range. Upper right: Cell-type accuracy (percentage of correctly predicted cells). Bottom left: Adjusted Rand Index (ARI). Evaluates the consistency of cell grouping structure and is insensitive to label imbalance or systematic label shifts. Bottom right: Normalized Mutual Information (NMI). Measures global correspondence between label distributions independent of exact label matches. **C)** Spatial evaluation of cell segmentation against Xenium reference. Left: Duplication rate. Quantifies segmentation granularity relative to Xenium, reflecting how many predicted cells map to the same reference cell. The metric is computed as the ratio of unique predicted cells to unique Xenium cells based on matched pairs, with values above 1 indicating over-segmentation. Right: Per-cell size Distribution. Cell size is approximated by the diagonal length of the bounding box enclosing each reconstructed cell. This metric provides a geometry-based proxy for comparing relative cell extents across segmentation methods. **D)** Gene expression level similarity between predicted cells and Xenium reference cells was evaluated on matched cell pairs using the top 2000 highly variable genes (HVGs). Top: Gene-level Spearman correlation, where higher values indicate stronger preservation of gene-wise rank structure. Middle: Gene-level RMSE, where lower values denote smaller expression deviations and improved quantitative agreement. Bottom: Gene-level Pearson correlation, where higher values reflect stronger linear correspondence of gene expression profiles.

To systematically evaluate STCS relative to existing methods, including Bin2Cell and 10x Space Ranger, we assessed three complementary levels of agreement with the ground truth based on CellTypist annotations.

At the cell level, predicted cells from each method were aligned to Xenium reference cells using maximum spatial overlap. This alignment enabled direct cell-wise comparisons with the ground truth. Across all evaluated metrics, including weighted per-class F1 score, cell-type accuracy, Adjusted Rand Index (ARI), and Normalized Mutual Information (NMI). STCS offers incremental improvement over existing methods (Fig. 2B).

At the spatial level, we examined mapping relationships between predicted and reference cells. Alignments were not always strictly one-to-one: single Xenium cells occasionally corresponded to multiple predicted cells, while multiple Xenium cells sometimes mapped to a single predicted cell. These events reflect differences in segmentation granularity. Consistent with its larger reconstructed cell sizes, 10x Space Ranger exhibited a higher duplication rate, multiple predicted cells mapping to the same Xenium cell, relative to Bin2Celll and STCS (Fig. 2C).

At the gene level, we evaluated similarity between predicted cells and their aligned Xenium cells using expression correlation and error metrics. STCS demonstrated marginal gains compared to existing methods across Spearman correlation, Pearson correlation, and RMSE (Fig. 2D).

These multi-level evaluations demonstrate that STCS achieves consistent improvements over existing segmentation methods across cell-type agreement, spatial granularity, and gene expression accuracy when benchmarked against Xenium ground truth, confirming its competitiveness as a cell segmentation approach for high-resolution spatial transcriptomics data

### STCS achieved superior spatial organization and transcriptomic accuracy on Stereo-seq data

To evaluate STCS on Stereo-seq data, we compared its performance against (i) STP, a Stereo-seq–specific single-cell partitioning framework (Li et al., Nature Communications, 2025), and (ii) a nuclei-only baseline based on StarDist nucleus segmentation. All methods were applied to the same Stereo-seq whole mouse brain dataset (17; 18). Cell-types were predicted using a CellTypist model, which provides fine-grained subtype annotations. Following the same crop-based parameter tuning strategy described above, we selected *S* = 30 and *λ* = 0.5 for this dataset. The larger search radius reflects the substantially finer spatial resolution of Stereo-seq bin1 units (0.5*µm ×* 0.5*µm*), which requires a broader neighborhood to capture comparable cellular extents. The resulting STCS segmentation and reconstructed cellular units with cell-class predictions are illustrated in 3A.

To facilitate robust spatial evaluation and reduce sensitivity to subtype-level noise, predicted labels were further consolidated into broader cell classes representing higher-level cellular categories. Because true single-cell ground-truth annotations are unavailable, performance metrics were evaluated by comparison to reference scRNA-seq data from whole mouse brain tissue (19).

Spatial organization was evaluated using metrics that quantify local identity consistency and spatial continuity. STCS demonstrated consistently higher cell-class coherence across neighborhood scales (Fig. 3B), indicating improved preservation of spatial identity structure and reduced local label discordance. In parallel, STCS achieved lower CHAOS scores at both the Leiden cluster and cell-class levels, reflecting decreased spatial disorder and clearer segregation of transcriptionally distinct populations. Lower CHAOS values indicate more spatially structured and biologically plausible organization of inferred cellular states.

**Figure 3:**
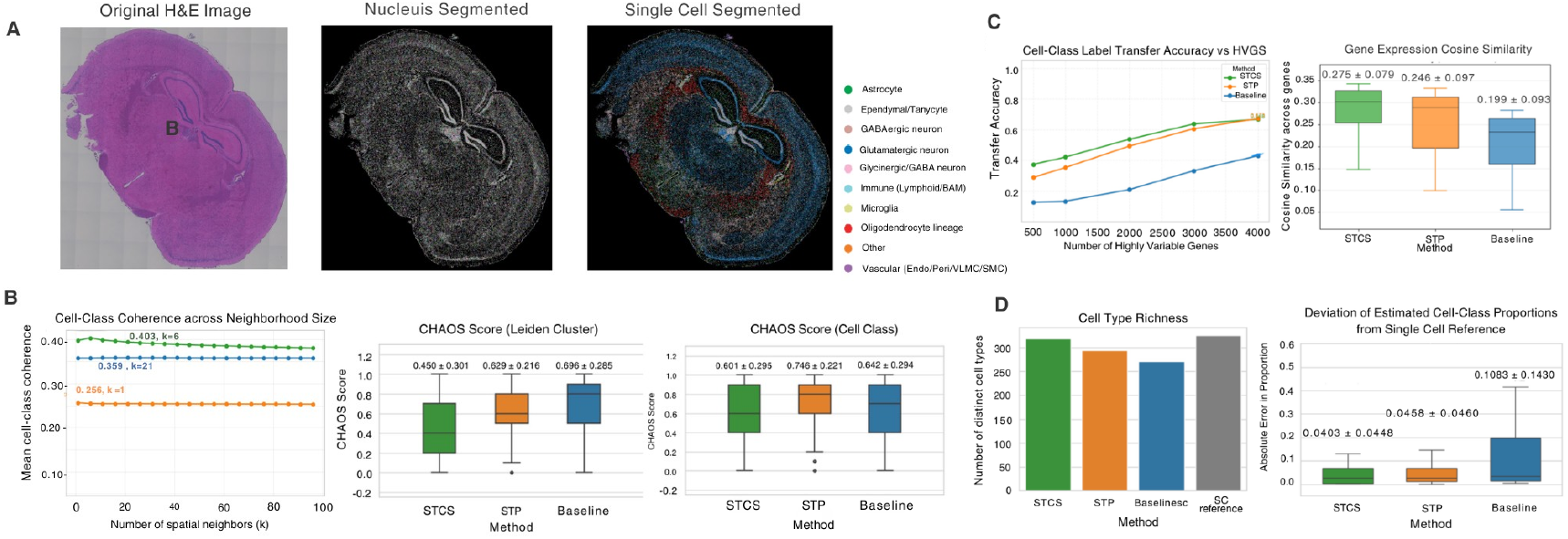
Performance of STCS on high-resolution Stereo-seq mouse brain data. **A)** Tissue images and segmentations. Left: original H&E image; Center: nuclei segmentation mask; Right: Single-cell Segmented slide by STCS (colored by cell class, defined as higher-level groupings of CellTypist-predicted cell types. **B)** Spatial continuity metrics. Left: Cell-class coherence across neighborhood size. computed as the proportion of spatially nearest neighbors sharing the same cell-class label at each neighborhood size. Higher values indicate greater spatial consistency of cell-class assignments within local tissue contexts. Middle: CHAOS scores computed at the Leiden cluster level. Right: CHAOS scores computed at the cell-class level. Lower CHAOS scores indicate better spatial organization. The center line represents the median CHAOS score and the box spans the interquartile range. The annotated text are *µ ± σ*. **C)** Gene Level metrics. Left: Cell-class label-transfer accuracy to a scRNA-seq reference across varying numbers of highly variable genes (HVGs), assessing how reliably reconstructed spatial cells recover reference-defined transcriptional identities and the robustness of predictions based on different selected features. Right: Distribution of gene-expression cosine similarity between reconstructed cells and scRNA-seq reference profiles across cell classes, quantifying transcriptomic agreement. Cosine similarity was computed between each reconstructed cell’s expression profile and the mean expression profile of its assigned cell class in the scRNA-seq reference. Center lines indicate medians and boxes span the interquartile range. The annotated text are *µ ± σ*. **D)** Cell-type composition accuracy. Left: Cell-type richness between methods and a single-cell reference data. Right: Absolute error in estimated cell-class proportions compared to the reference. Lower values indicate closer agreement with the cellular composition of the reference data. The annotated text are *µ ± σ* and center lines indicate median.

At the gene level, reconstructed profiles were evaluated against the scRNA-seq reference using label-transfer accuracy and cosine similarity computed within cell classes (Fig. 3C). STCS yielded higher values on both metrics, indicating that aggregated transcripts within STCS-segmented cells more closely align with reference-defined transcriptional states and exhibit greater molecular similarity to their corresponding reference populations.

Besides the high spatial coherence and gene-expression similarity to the reference data, STCS exhibited the highest cell-type richness among all evaluated methods (Fig. 3D). Additionally, STCS achieved the closest agreement with the reference cell-class proportions, as reflected by the lowest absolute error in the proportions compared to the reference data.

Across all evaluated dimensions, including spatial organization, gene-expression accuracy, and cell-type composition, STCS consistently outperformed both STP and the nuclei-only baseline, demonstrating its effectiveness as a cell segmentation method for high-resolution Stereo-seq data without requiring ground-truth annotations.

## Discussion

In summary, STCS addresses the central challenge of reconstructing coherent cell-level units from sequencing-based spatial transcriptomics data, where measurements are inherently defined over spatial bins rather than biological cells. Across datasets and evaluation settings, STCS achieves robust performance in cell segmentation, preserving spatial structure while maintaining biologically consistent transcriptional profiles. Unlike most current approaches, which are primarily restricted to 10x Genomics platforms, STCS is designed for broad compatibility with sequencing-based spatial transcriptomics technologies. We demonstrate its applicability on both Visium HD and Stereo-seq datasets, with consistent performance across platforms, and provide STCS as an open-source framework to support reproducible and extensible use.

STCS offers flexibility in accommodating slides from different tissue types through adjustable parameters that adapt to variations in tissue composition and spatial resolution. We provide reference-free evaluation metrics, including transcript capture saturation, cell-type stability, and spatial coherence, to guide parameter selection without requiring ground-truth annotations. However, in the absence of matched ground-truth data such as Xenium, comprehensive performance evaluation remains challenging. In future work, we plan to explore learning-based approaches for automated parameter selection across tissue types, which may further enhance the robustness of STCS and enable recommended parameter configurations for diverse experimental settings.

Furthermore, we established an evaluation pipeline using slides that contain both Xenium and Visium HD data, enabling direct cell-level benchmarking against Xenium-derived annotations. However, such datasets are generated using post-Xenium Visium HD protocols and present several limitations. Xenium captures a targeted gene panel rather than the transcriptome-wide coverage of Visium HD, restricting the scope of expression-level comparisons (10; 6). Additionally, Visium HD data acquired after Xenium processing may exhibit reduced data quality due to sequential tissue handling (20). Moreover, the reliability of Xenium cell segmentation itself varies across cells. Xenium employs a multimodal segmentation pipeline that prioritizes boundary staining using surface markers, interior RNA staining, and nucleus expansion in decreasing order of expected accuracy (9). In the lung cancer dataset used in this study, only 15.8% of cells were segmented by boundary staining, while 82.0% relied on interior RNA staining and 2.2% on nucleus expansion alone (21), introducing uncertainty in cell boundary placement that may propagate into downstream benchmarking comparisons (22). The limited availability of dual-modality slides further constrains comprehensive evaluation. Together, these factors underscore the need for continued refinement of this benchmarking strategy and its validation across a broader range of samples and experimental settings (6).

## Acknowledgements

We thank past and present members of the Yang lab for valuable scientific discussions. This work was supported by start-up funding from the School of Arts and Sciences and the Human Genetics Institute of New Jersey at Rutgers University (J.Y.), as well as seed funding from the School of Arts and Sciences (J.Y. and L.L.).

## Author Contributions

This study was designed and directed by J.Y., and L.L. L.C.W., and X.H. developed the algorithm. L.C.W., X.H., F.Z., C.S., and J.G. performed data analysis. R.O., T.T., and M.P.V. provided scientific input. L.C.W., X.H., and J.Y. wrote the manuscript.

## Methods

### STCS Pipeline

#### Nuclei Segmentation and Initialization

STCS takes as input a paired histological (H&E) image *I* and a spatial transcriptomics dataset consisting of *N* spatially barcoded bins. Each bin *i* is characterized by its spatial coordinates *s*_*i*_ ∈ ℝ ^2^ and a gene expression vector *x*_*i*_ ∈ ℝ^*G*^, where *G* is the number of genes.

First, STCS applies StarDist, a deep learning-based nuclear segmentation model, to the H&E image to identify *C* cell nuclei. This produces a set of nucleus masks 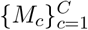, where *M*_*c*_ ⊂ ℝ^2^ denotes the set of pixel coordinates belonging to nucleus *c*.

Each bin is then assigned an initial nucleus label based on whether its spatial coordinates fall within a detected nucleus:

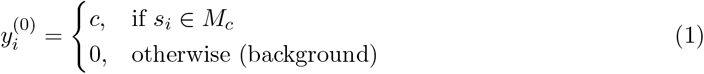

#### Nucleus-level Pseudobulk Construction

For nuclei with at least one assigned bin, STCS constructs nucleus-level expression profiles by aggregating associated bins:

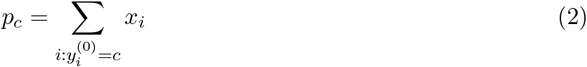

where *x*_*i*_ ∈ ℝ^*G*^ is the expression vector of bin *i*. This yields a nucleus-by-gene matrix *P* ∈ ℝ^*C×G*^.

#### Feature Preprocessing and Latent Space Projection

The nucleus-level expression profiles *p*_*c*_ are preprocessed using gene filtering, normalization, log transformation, feature selection, and scaling. Let 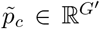 denote the preprocessed profile for nucleus *c*, where *G*^*′*^ is the number of retained genes. Principal component analysis (PCA) is then applied to obtain low-dimensional representations:

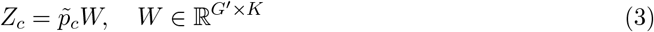

where *W* is the PCA projection matrix and *K* is the number of principal components.

Each bin undergoes identical preprocessing, yielding 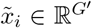, and is projected into the same latent space:

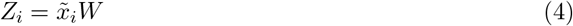

If a single-cell RNA-seq reference is provided, STCS computes a weighting matrix Σ ∈ ℝ^*K×K*^ as the pairwise cosine similarity between principal components evaluated across reference cells, which is used to weight the transcriptomic distance.

### Spatial Candidate Nuclei Selection

For each bin *i*, STCS restricts assignment to a set of candidate nuclei defined by spatial proximity:

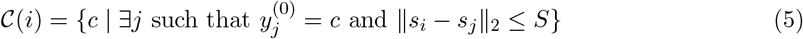

where *S* is the search radius parameter.

### Joint Transcriptomic–Spatial Assignment

For each bin–nucleus pair (*i, c*) with *c* ∈ *𝒞* (*i*), STCS computes a transcriptomic distance:

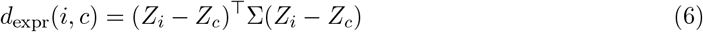

where Σ is the similarity-derived weighting matrix in latent space. Spatial distance is defined as:

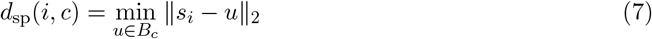

with *B*_*c*_ denoting the set of bin locations initially assigned to nucleus *c*. The normalized transcriptomic distance 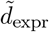 (*i, c*) is obtained by applying a log transformation followed by min–max normalization to the raw transcriptomic distances across all candidate pairs:

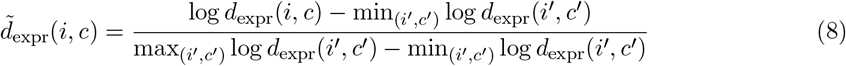

The combined assignment score is:

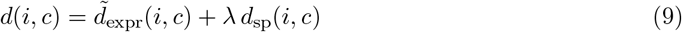

where *λ* controls the spatial weighting. Each bin is assigned to the nucleus minimizing this score:

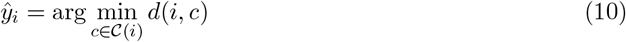

### Cell-level Reconstruction and Annotation

Final cell-level expression profiles are reconstructed by aggregating all bins assigned to each nucleus:

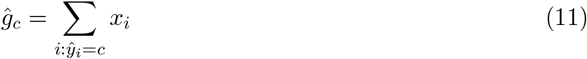

The resulting assignments define inferred cell boundaries and corresponding cell-resolved gene expression profiles. Cell-type annotation is then performed using CellTypist or other cell type annotation method on the reconstructed cell-level profiles.

### Downstream analysis

#### Detected Genes per Cell

To assess the transcriptomic completeness of reconstructed cells across parameter settings, we computed the mean number of detected genes per cell. For each cell *c*, the number of detected genes is defined as:

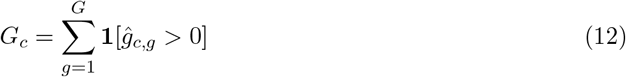

where 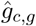 is the reconstructed expression of gene *g* in cell *c*. To ensure comparability across parameter combinations, the metric was evaluated on a shared set of cells defined as the intersection of cell identities across all parameter configurations within each crop. The mean detected-gene count was then averaged across crops, with standard deviation reported as error bars.

#### Cell Counts per Cell Type

To evaluate the stability of cell-type predictions across parameter settings, we quantified the number of cells assigned to each cell type as a function of the search radius *S*. Cell-type labels were obtained by applying CellTypist to the reconstructed cell-level expression profiles at each parameter configuration. For each cell type *k* and search radius *S*, the cell count is:

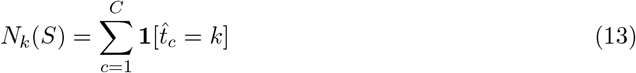

where 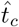 denotes the predicted cell type for cell *c* and *C* is the total number of reconstructed cells. Stable cell counts across increasing *S* indicate that cell-type composition is robust to changes in the search radius.

#### Connection Score

To quantify the spatial coherence of reconstructed cells, we define a connection score that measures the proportion of spatial units forming a contiguous region within each cell. For a given cell *c*, let *B*_*c*_ = *{b*_1_, *b*_2_, …, *b*_*n*_*}* denote the set of spatial coordinates assigned to that cell. An adjacency graph *G*_*c*_ = (*B*_*c*_, *E*_*c*_) is constructed, where an edge (*b*_*j*_, *b*_*k*_) ∈ *E*_*c*_ exists if and only if *b*_*j*_ and *b*_*k*_ are spatially adjacent under an 8-connected neighborhood (i.e., horizontal, vertical, and diagonal neighbors on the grid). The connection score for cell *c* is then defined as:

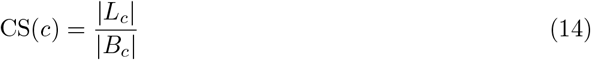

where *L*_*c*_ is the largest connected component of *G*_*c*_. Values approaching 1 indicate that nearly all spatial units assigned to the cell form a single contiguous region, reflecting spatially coherent segmentation. Lower values indicate fragmented assignments in which spatial units belonging to the same cell are spatially disconnected.

#### Parameter Sensitivity Analysis

To characterize the sensitivity of STCS performance to the search radius *S* and spatial weighting factor *λ*, we evaluated cell-type agreement and compositional divergence across a grid of parameter combinations. For each combination (*S, λ*), STCS was applied independently to five randomly sampled crops, and matched cell pairs were constructed by aligning predicted cells to Xenium reference cells via spatial overlap.

Cell-type agreement was quantified using the Adjusted Rand Index (ARI), computed between predicted and reference cell-type labels on matched pairs. Compositional divergence was assessed using the Jensen–Shannon divergence (JSD) between the predicted and reference cell-type proportion vectors.

To isolate the effect of each parameter, we computed marginal summaries by averaging over the complementary parameter and across all crops. Specifically, the effect of *S* was obtained by averaging ARI and JSD across all *λ* values and crops for each fixed *S*:

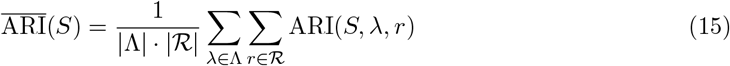

where Λ denotes the set of evaluated *λ* values and *ℛ* denotes the set of crops. The effect of *λ* was computed analogously by averaging over all *S* values and crops. Standard deviations across the averaged configurations were reported as error bars. Optimal parameter values were identified as those maximizing ARI (or minimizing JSD) in the marginal summaries.

#### Cell-level Alignment

To enable direct cell-level comparisons between reconstructed cells and Xenium reference cells, we established one-to-one cell correspondences through a three-stage alignment procedure.

##### Stage 1: Candidate identification via nucleus spatial join

For each predicted cell *c*_pred_, the StarDist nucleus segmentation provides a set of nucleus pixel coordinates. Xenium reference cells are represented as polygons 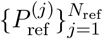, constructed from the convex hull of all spatial units assigned to each Xenium cell. Each nucleus point **n**_*i*_ associated with a predicted cell is assigned to the Xenium polygon that contains it:

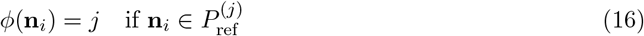

When a predicted cell contains multiple nucleus points mapping to different Xenium polygons, the candidate reference cell is determined by majority vote:

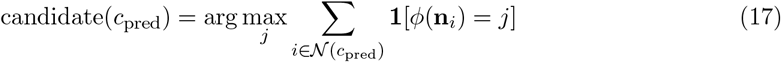

where *𝒩* (*c*_pred_) denotes the set of nucleus points belonging to predicted cell *c*_pred_. This stage produces a set of candidate predicted–reference cell pairs.

##### Stage 2: Boundary refinement via polygon intersection-over-union

To verify and refine the candidate alignments, we assessed spatial boundary agreement between matched cells using the intersection-over-union (IoU). For each candidate pair, cell boundaries were approximated by constructing alpha-shape polygons (*α* = 0.05) from the spatial coordinates of all spatial units assigned to each cell. The IoU between predicted polygon *P*_pred_ and reference polygon *P*_ref_ is defined as:

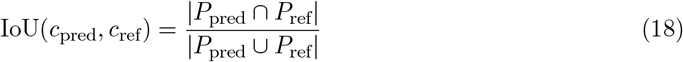

where | · | denotes polygon area. To resolve ambiguous candidates, bidirectional matching was performed based on maximum intersection area. For each reference polygon, the predicted polygon with the largest intersection area was selected as the forward match:

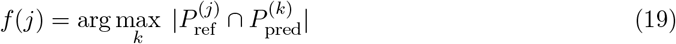

The reverse match *r*(*k*) was computed analogously. Candidate polygons were efficiently identified using a spatial R-tree index to restrict intersection tests to geometrically overlapping pairs.

##### Stage 3: One-to-one matching

Final matched pairs were retained only when the forward and reverse matches agreed reciprocally, i.e., *f* (*j*) = *k* and *r*(*k*) = *j*. Pairs that did not satisfy this reciprocal criterion were excluded. Additionally, duplicate mappings were removed on both sides to ensure that each predicted cell maps to exactly one reference cell and vice versa. The resulting set of one-to-one matched pairs 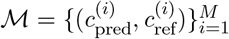 was used for all subsequent cell-level evaluations, including cell-type agreement, gene-expression comparison, and spatial metric computation.

### Cell-type Agreement Metrics

Given a set of matched cell pairs with ground-truth labels 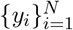 and predicted labels 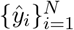, we evaluated cell-type agreement using the following metrics.

*Cell-type accuracy* is defined as the proportion of matched cells for which the predicted cell type equals the ground-truth label:

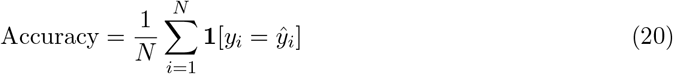

*Adjusted Rand Index* (ARI) measures the agreement between two label partitions, corrected for chance. Given a contingency table between ground-truth and predicted labels, ARI is computed as:

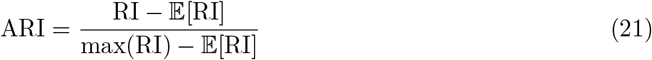

where RI is the Rand Index. Values near 1 indicate strong agreement, while values near 0 indicate chance-level correspondence.

*Normalized Mutual Information* (NMI) quantifies the mutual dependence between the groundtruth and predicted label distributions, normalized to the range [0, 1]:

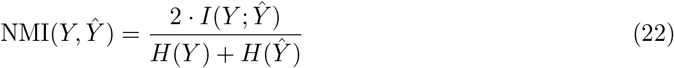

where *I*(*Y* ; Ŷ) is the mutual information and *H*(·) denotes entropy. Higher values indicate stronger global correspondence between label distributions.

### Per-class and Weighted F1 Score

For each cell type *k*, the F1 score is computed as the harmonic mean of precision and recall:

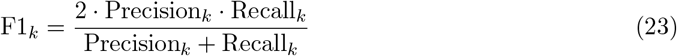

The weighted F1 score aggregates per-class F1 values weighted by class support (number of matched cells per class):

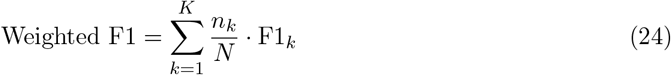

where *n*_*k*_ is the number of matched cells belonging to class *k* and *N* = Σ_*k*_ *n*_*k*_.

### Jensen–Shannon Divergence

To assess agreement between predicted and reference cell-type compositions at the population level, we computed the Jensen–Shannon divergence (JSD) between the two cell-type proportion distributions. Let *p* and *q* denote the normalized cell-type frequency vectors derived from groundtruth and predicted annotations, respectively. The JSD is defined as:

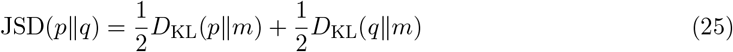

where 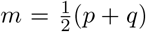 and *D*_KL_ denotes the Kullback–Leibler divergence. JSD is bounded in [0, 1] (when using base-2 logarithm), with lower values indicating closer agreement between the two distributions.

### Duplication Rate

To assess segmentation granularity relative to the Xenium reference, we computed the duplication rate for each method. Predicted cells were aligned to Xenium reference cells based on nuclear spatial overlap, and the duplication rate was defined as the ratio of unique predicted cells to unique reference cells among all matched pairs:

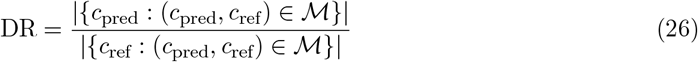

where *ℳ* denotes the set of matched predicted–reference cell pairs. Values above 1 indicate over-segmentation, where multiple predicted cells map to a single reference cell. Values below 1 indicate under-segmentation.

### Per-cell Size Distribution

To compare the spatial extent of reconstructed cells across methods, we computed per-cell size as the diagonal length of the bounding box enclosing all spatial units assigned to each cell. For a cell *c* with assigned spatial coordinates *{s*_*j*_*}*_*j*∈*Ac*_, the bounding box width and height are:

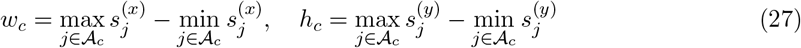

and the cell size is defined as:

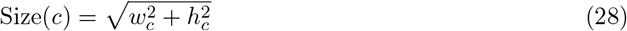

This metric provides a geometry-based proxy for comparing relative cell extents across segmentation methods without requiring ground-truth cell boundaries.

### Gene-level Expression Metrics

Gene-level agreement between reconstructed and reference cells was evaluated on matched cell pairs using the top 2,000 highly variable genes (HVGs) identified from the Xenium reference. HVGs were selected using the Seurat v3 method applied to the subset of genes shared between the predicted and reference datasets.

For each matched pair (*c*_ref_, *c*_pred_), let **x**, 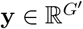 denote their expression vectors over the selected HVGs. A both-nonzero mask *ℳ* = *{g* : *x*_*g*_ ≠ 0 and *y*_*g*_ ≠ 0*}* was applied to restrict comparisons to genes with detectable expression in both cells. Let **x**_*M*_, **y**_*M*_ denote the masked vectors restricted to *ℳ*, with |*ℳ*| ≥ 2 required for metric computation.

*Pearson correlation* measures the linear correspondence between matched expression profiles:

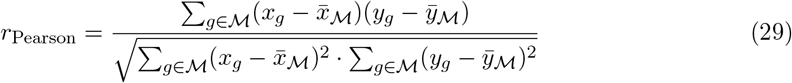

*Spearman correlation* measures the rank-order agreement between expression profiles. Gene expression values within ℳ are replaced by their ranks, and the Pearson correlation is computed on the ranked vectors:

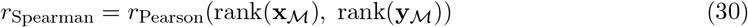

*Root mean squared error* (RMSE) quantifies the magnitude of expression deviations between matched cells:

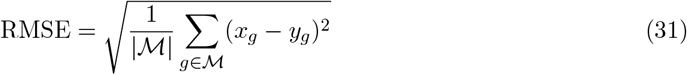

### Per-gene Expression Agreement

In addition to per-cell metrics, we evaluated expression agreement at the gene level across all matched pairs. For each gene *g* among the selected HVGs, the Pearson correlation was computed across matched cells using a streaming algorithm:

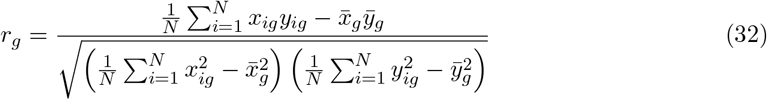

where *x*_*ig*_ and *y*_*ig*_ denote the expression of gene *g* in the *i*-th reference and predicted cell, respectively, and *N* is the number of matched pairs. Higher per-gene correlations indicate that the reconstructed expression faithfully recapitulates cell-to-cell variation observed in the reference for that gene.

### CHAOS Score

To quantify the spatial organization of inferred clusters and cell classes, we computed the CHAOS score, which measures the proportion of spatially adjacent cells assigned to different labels. For each cell *c* with label *ℓ*_*c*_, the *k* nearest spatial neighbors are identified using Euclidean distance on spatial coordinates. The per-cell CHAOS score is defined as:

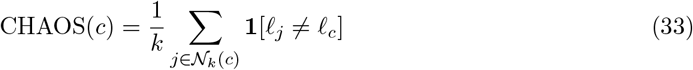

where 𝒩_*k*_(*c*) denotes the set of *k* spatially nearest neighbors of cell *c* (excluding *c* itself), and *k* = 6 was used throughout. Lower CHAOS scores indicate greater spatial coherence, meaning cells sharing the same label tend to co-localize in tissue space. The metric was computed separately using Leiden cluster labels and cell-class labels to assess spatial organization at different levels of annotation granularity.

### Cell-class Coherence

Cell-class coherence quantifies the degree to which spatially neighboring cells share the same cell-class identity. For each cell *c* with cell-class label *t*_*c*_, the coherence score at neighborhood size *k* is defined as:

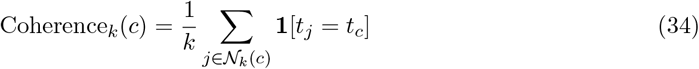

where 𝒩_*k*_(*c*) denotes the *k* spatially nearest neighbors of cell *c*. The mean coherence across all cells provides a global measure of spatial identity consistency:

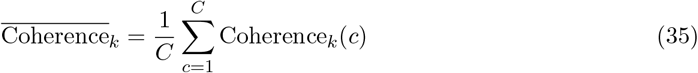

Higher values indicate that cells of the same class are spatially clustered. To evaluate robustness across spatial scales, coherence was computed over a range of neighborhood sizes *k* ∈ *{*1, 6, 11, …, 96*}*, and the resulting profiles were compared across methods.

### Label-transfer Accuracy

To evaluate how well reconstructed spatial cells recover reference-defined transcriptional identities, we performed label transfer from the spatial dataset to a held-out scRNA-seq reference. For each scRNA-seq cell, a *k*-nearest neighbor search (*k* = 3) was performed against the spatial dataset in a shared highly variable gene (HVG) space. Expression vectors were L2-normalized and cosine distance was used as the similarity metric. The transferred label for each scRNA-seq cell was determined by inverse-distance-weighted majority voting among its *k* spatial neighbors:

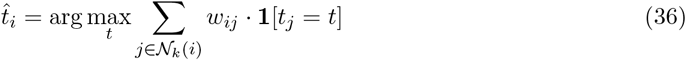

where 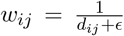 is the inverse-distance weight, *d*_*ij*_ is the cosine distance between scRNA-seq cell *i* and spatial neighbor *j*, and *t*_*j*_ is the cell-class label of spatial cell *j*. Transfer accuracy was computed as the proportion of scRNA-seq cells for which the transferred label matched the ground-truth cell-class annotation:

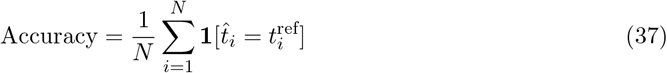

To assess robustness to feature selection, the evaluation was repeated across varying numbers of HVGs (500, 1,000, 2,000, 3,000, 4,000), where HVGs were selected from the scRNA-seq reference using the Seurat method. Only genes present in both the spatial and reference datasets were considered.

### Gene-expression Cosine Similarity to Reference

To quantify the transcriptomic agreement between reconstructed spatial cells and the scRNA-seq reference at the cell-class level, we computed the cosine similarity between each reconstructed cell’s expression profile and the mean expression profile of its assigned cell class in the reference. For a reconstructed cell *c* assigned to cell class *k*, let 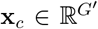 denote its expression vector over the selected HVGs and 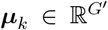 denote the mean expression vector of class *k* in the scRNA-seq reference. The cosine similarity is:

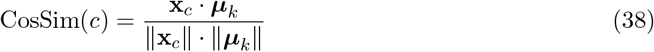

Higher values indicate that the reconstructed cell’s expression profile more closely resembles the reference transcriptional state of its assigned class. The distribution of cosine similarities was summarized per cell class and compared across methods using box plots, with median, interquartile range, and annotated mean *±* standard deviation reported.

### Cell-type Richness

To assess the diversity of cell-type predictions, we computed the cell-type richness for each method, defined as the number of distinct cell types identified:

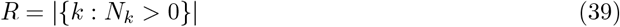

where *N*_*k*_ is the number of cells assigned to cell type *k*. Richness was compared against the scRNA-seq reference to evaluate whether each method recovers the expected cellular diversity. We additionally reported coverage, defined as the fraction of reference cell types recovered by each method:

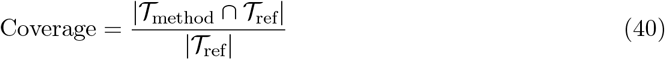

where *𝒯*_method_ and *𝒯*_ref_ denote the sets of cell types identified by the method and reference, respectively.

### Cell-type Proportion Error

To quantify how closely reconstructed cell-type compositions match the scRNA-seq reference, we computed the per-cell-type absolute error in estimated proportions. Let *p*_*k*_ and *q*_*k*_ denote the proportion of cells assigned to cell type *k* in the reference and predicted datasets, respectively. The absolute error for cell type *k* is:

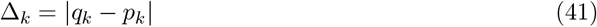

The distribution of Δ_*k*_ across all cell types was summarized using box plots, with lower values indicating closer agreement with the reference composition. Proportions were computed over the union of cell types across all methods and the reference, with missing types assigned a proportion of zero.

## Data availability

The Visium HD human lung cancer dataset with matched Xenium annotations used for ground-truth benchmarking is publicly available from 10x Genomics. The Visium HD are available at:(https://www.10xgenomics.com/datasets/visium-hd-cytassist-gene-expression-human-lung-cancer-post-xenium-expt), and the corresponding Xenium data at: (https://www.10xgenomics.com/datasets/xenium-human-lung-cancer-post-xenium-technote). The single-cell RNA-seq reference used for CellTypist model training was obtained from (16).

The Stereo-seq whole mouse brain dataset is available from STOmics (https://en.stomics.tech/col1241/index.html). The whole mouse brain single-cell reference used for Stereo-seq evaluation is available from the Allen Brain Cell Atlas (https://mouse.brain-map.org/static/atlas).

## Code availability

The code is released on the github: https://github.com/YangLabRutgers/STCS.

